# Inoculum-dependent bactericidal activity of a *Mycobacterium tuberculosis* MmpL3 inhibitor

**DOI:** 10.1101/2023.02.22.529622

**Authors:** Bryan Berube, Aditi Deshpande, Amala Bhagwat, Tanya Parish

## Abstract

Indolcarboxamides are a promising series of anti-tubercular agents which target *Mycobacterium tuberculosis* MmpL3, the exporter of trehalose monomycolate, a key cell wall component. We determined the kill kinetics of the lead indolcarboxamide NITD-349 and determined that while kill was rapid against low density cultures, bactericidal activity was inoculum-dependent. A combination of NITD-349 with isoniazid (which inhibits mycolate synthesis) had an increased kill rate; this combination prevented the appearance of resistant mutants, even at higher inocula.

---

*Mycobacterium tuberculosis* remains a pathogen of global significance, causing millions of new cases of tuberculosis and ∼1.6 million deaths in 2022 (1). A substantial effort in identifying new anti-tubercular drugs has been made in the last decade and several novel agents are in or approaching clinical trials. Cell wall biosynthesis is a major target of new agents, with many novel chemical series inhibiting the action of the mycolic acid exporter MmpL3 (2). The indolcarboxamide series has been the subject of many studies since it has demonstrated excellent potency against extracellular *M. tuberculosis* and *in vivo* efficacy in animal models of infection (3).

Previous work has demonstrated that the indolcarboxamide NITD-304 is a potent inhibitor of bacterial growth with bactericidal activity. However, kill kinetics were moderate with a 1-log reduction in viable bacteria at the IC_50_ and 4-log of kill at 100X IC_50_ without complete sterilization of the culture (3). We were interested to determine if this was a property of the series or the molecule. Therefore we looked at the kill kinetics of NITD-349, the most advanced molecule in the series. We determined the minimum inhibitory concentration (MIC) using our standard growth assay (4); bacteria were inoculated at an OD of 0.02 and incubated for 5 days – the IC_50_ was 56 ± 22 nM and the IC_90_ was 84 ± 17 n,M(n=4) which is in line with the reported IC_50_ of 23 nM (3).

We determined the minimum bactericidal concentration (MBC), defined as the lowest concentration giving 3-log kill within 21 days (5). We measured viability by colony forming units after exposure to NITD-349; bacteria were inoculated into Middlebrook 7H9 medium plus 10% v/v Middlebrook OADC supplement (oleic acid, albumin, dextrose, catalase) and 0.05% w/v Tween 80. Medium contained NITD-349 prepared from a 10 mM stock in DMSO; all cultures had DMSO to a final concentration of 1%. CFUs were measured by plating ten-fold serial dilutions onto Middlebrook 7H10 medium plus 10% v/v OADC and incubating for 3-4 weeks at 37°C.

We determined that NITD-349 was rapidly bactericidal with a complete sterilization of culture within 7 days at 125 nM (MBC = 125 nM) (Figure 1A). We repeated the kill kinetic study and saw a similar profile with rapid kill within 7 days and an MBC of 125 nM (Figure 1B). However in the second run we saw outgrowth at 125 nM after 7 days, presumably due to the growth of resistant mutants at the lower concentrations. We ran a third kill kinetic study with additional time points and demonstrated that kill was extremely rapid, with an almost linear kill over the first 7 days and all concentrations demonstrating complete sterilization within 14 days (MBC = 63 nM) (Figure 1C). Given the rapid kill rate, there was little scope to determine if kill was concentration-dependent or time-dependent. Of interest to us was the fact we saw much higher kill rates than the initial study; a comparison of the two studies showed that we used a lower inoculum starting at <10^6^ CFU/mL as compared to 10^7^ CFU/mL for NITD-304 (3).

**Figure 1.**
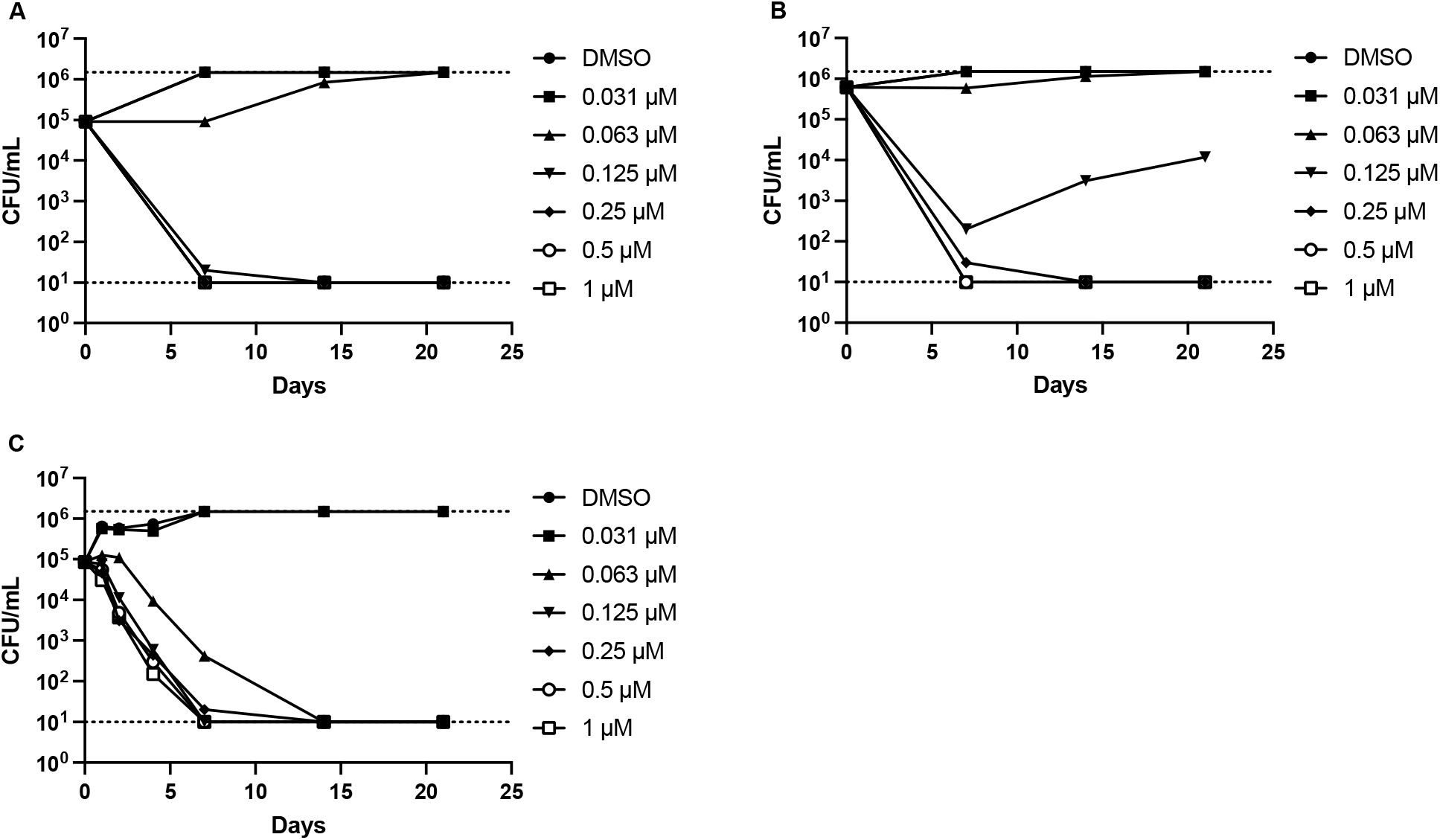
Kill kinetics of NITD-349. *M. tuberculosis* was inoculated into medium containing NITD-349 to a theoretical OD_600_ of 0.001 (A and C) or 0.005 (B). Viable bacteria were enumerated by plating serial dilutions and counting CFUs after 3-4 weeks. The upper and lower limits of detection are indicated by dotted lines. Day 0 values are for the DMSO control.

We repeated our kill kinetics studies using different inocula ranging from ∼10^5^ to ∼10^7^ CFU/mL. Each experiment was repeated three times in fully independent runs. We saw a clear inoculum-dependent effect (Figure 2). At the lowest inoculum we obtained 3 log kill with an MBC of 125 nM as in our prior experiment (Figure 2A). As we increased the inoculum, the kill rate decreased; at both higher inocula the MBC was >1 μM (Figure 2B and 2C). At a starting inoculum of >10^6^ the kill rate was 1.5 log per week, although the MBC was >1 μM; the highest concentration did lead to a 2.5 log kill. Kill was seen with concentrations as low as 125 nM (Figure 2B). At a starting inoculum of >10^7^, much less kill was apparent, with a maximum kill rate of 1 log per week and the highest concentration led to a 2 log kill. Kill was seen at concentrations about 250 nM (Figure 2C). This shows a clear inoculum dependent effect, such that kill rates decrease as the starting inoculum is higher and likely explains the difference in kill rates between NITD-304 and NITD-349.

**Figure 2.**
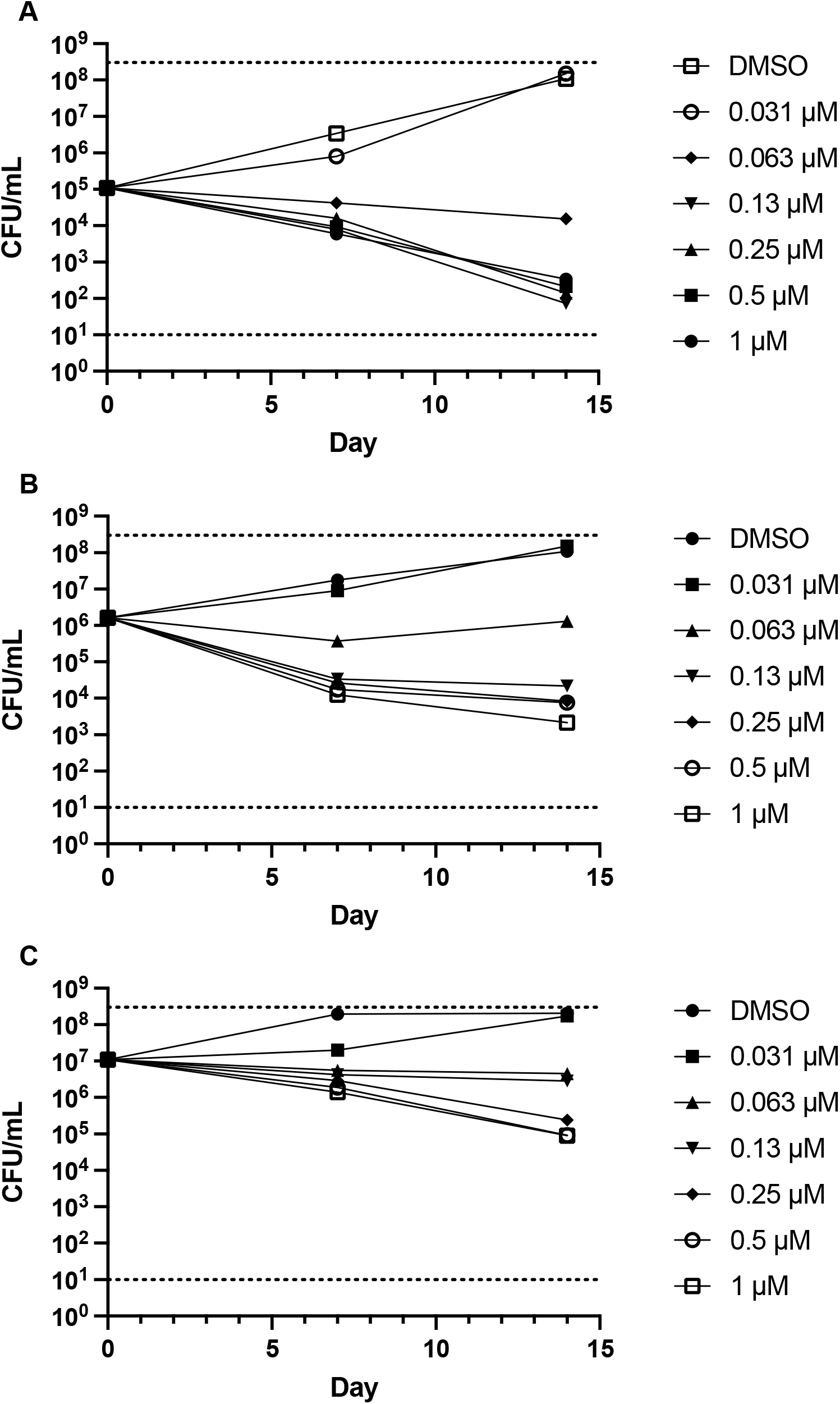
Inoculum-dependent kill kinetics of NITD-349. *M. tuberculosis* was inoculated into medium containing NITD-349 to a theoretical OD_600_ of (A) 0.001, (B) 0.01, and (C) 0.1. Viable bacteria were enumerated by plating serial dilutions and counting CFUs after 3-4 weeks. Data are the mean and standard deviation of three independent experiments. The upper and lower limits of detection are indicated by dotted lines. Day 0 values are for the DMSO control.

TB therapy consists of multiple drugs as a regimen, so it is important to know how different agents combine. Isoniazid is one of the frontline drugs for tuberculosis, but the frequency of resistance *in vitro* is high since it is a pro-drug. INH targets mycolic acid synthesis and so should synergize with MmpL3 inhibitors like NITD-349. We determined the effect of combining NITD-349 and INH on kill kinetics and the inoculum-dependent effect (Figure 3). We started with a low inoculum for this experiment since the frequency of resistance to INH is high (about 1 in 10^6^). NITD-349 showed rapid kill over the first 7 days as expected, with sterilization of the culture by Day 14. Addition of INH to NITD-349 increased the rate of kill by ∼1 log over the first 7 days, although INH alone was insufficient to kill.

**Figure 3.**
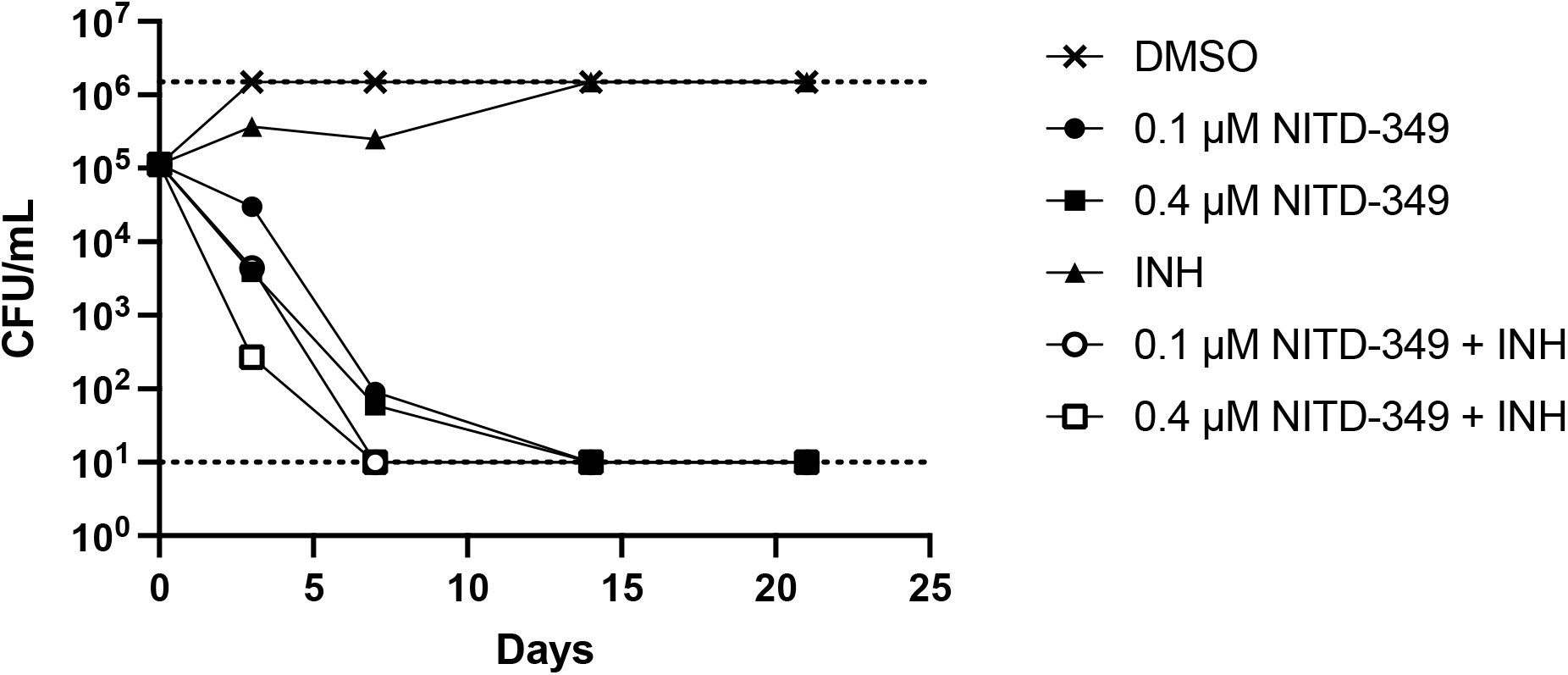
Combination kill kinetics of NITD-349 and INH. *M. tuberculosis* was inoculated into medium containing NITD-349 and/or 5 μM INH to a theoretical OD_600_ of 0.001. Viable bacteria were enumerated by plating serial dilutions and counting CFUs after 3-4 weeks. The upper and lower limits of detection are indicated by dotted lines. Day 0 values are for the DMSO control.

We repeated the experiment as two fully independent experimental runs using different inocula to determine whether we still saw the inoculum-dependent effect in combinations (Figure 4). Since the appearance of resistant mutants is random, the number of resistant bacteria in each starting culture will be different, therefore we present the data for each experiment separately. At the lowest inoculum we saw rapid kill by NITD-349 and INH alone as expected (Figure 4A to 4C). In contrast to our previous run, we did see kill by INH, which is likely due to difference in the number of INH resistant bacteria in the inoculum. At the lower inocula, we saw outgrowth of resistant mutants with INH in both experiments and for NITD-349 in one experiment at the lower concentration. Combination of the two agents led to improved kill rates of at least 1 log over 7 days and prevented the appearance of resistant mutants (Figure 4A and 4C). Using a slightly larger inoculum, we saw outgrowth of resistant mutants for INH and NITD-349 at the lowest concentration in both experiments; again the combination improved kill rates and prevented the appearance of resistant mutants (Figure 4B and 4D). Using the highest inoculum, we saw limited kill with NITD-349 alone confirming our previous observation (Figure 4C and 4E). In each of the runs, we saw early kill by INH followed by outgrowth of resistant mutants. However, in combination we saw an improved kill rate and no appearance of resistant mutants. Thus we conclude that the combination of NITD-349 and INH can prevent the appearance of resistant mutants. Although kill rates were reduced for the higher inoculum, they were improved in the combination treated cultures, therefore the combination of two agents can overcome the inoculum-dependent effect to some extent.

**Figure 4.**
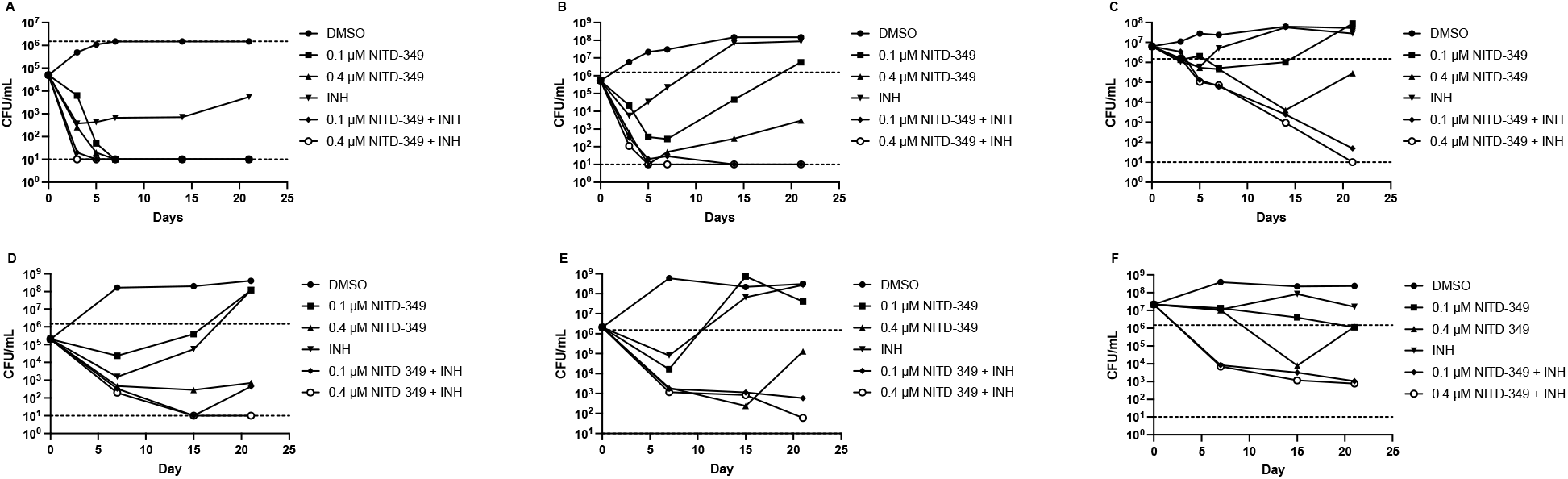
Inoculum-dependent combination kill kinetics of NITD-349 and INH. *M. tuberculosis* was inoculated into medium containing NITD-349 and/or 5 μM INH to a theoretical OD_600_ of 0.001 (A and D), 0.01 B and E) and 0.1 (C and F). Viable bacteria were enumerated by plating serial dilutions and counting CFUs after 3-4 weeks. Two independent experiments were conducted in panels A-C and D-F. The upper and lower limits of detection are indicated by dotted lines. Day 0 values are for the DMSO control.

In conclusion, we have shown that the bactericidal activity of NITD-349 is inoculum-dependent, but this can be mitigated by combination with INH. The combination of NITD-349 and INH showed increased kill rates and prevented the appearance of resistant mutants, suggesting that this combination could be clinically useful.

## Acknowledgements

We thank Kyle Krieger for technical assistance in performing kill kinetics. We thank Anna Upton, Khisi Mdluli and Nader Fotouhi at the TB Alliance for supplying NITD-349 and useful discussion.

